# Assessment of the Effects of Environmental Perturbations on Soil Ecology in A Terrestrial Mesocosm

**DOI:** 10.1101/2023.07.21.550096

**Authors:** Kathleen L. Arnolds, Riley C. Higgins, Jennifer Crandall, Gabriella Li, Jeffrey G. Linger, Michael T. Guarnieri

**Author notes:** **Corresponding Author:** Michael T. Guarnieri, Ph.D. 15013 Denver West Pkwy Golden, CO, USA 80401.

## Abstract

Climate change is altering ecosystems in unprecedented ways and necessitates the development of strategies that model ecosystems and allow for the evaluation of environmental impacts of perturbations: including climate events, novel approaches to agronomy or ecosystem management, and impacts of bio-industry and biotechnology innovations. Mesocosms present a platform to model some of the complexity of an ecosystem, while still being controlled and reproducible enough that they can be used to ask targeted questions and systematically assess the impacts of perturbation events. Herein, we established a methodological pipeline to assess the impact of three perturbation events (hydration, nutrification, contamination) upon plant-associated microbial communities using a terrestrial mesocosm. Mesocosms were assessed over a 30-day time-course following environmental perturbations, including modeling contamination with a foreign microbe via the introduction of *Saccharomyces cerevisiae*. We developed and applied a suite of diagnostic and bioinformatic analyses, including digital droplet PCR, microscopy, and phylogenomic analyses to assess the impacts of a perturbation event in a system that models a terrestrial ecosystem. The resultant data show that our mesocosms are dynamic yet reproducible, and that the analysis pipeline presented here allowed for a longitudinal assessment of microbial population dynamics and abiotic soil characteristics following perturbations, as well as the fate of yeast in the soil. Notably, our data indicate that a single perturbation event can have long-lasting impact upon soil composition and underlying microbial populations. Thus, this approach can be used to ask targeted questions as well as gain insights on broader ecological trends of soil perturbation events.

**Importance:** Soils are key to a healthy environment, but the impact of human activities and climate change upon soil microbiomes remains unclear. It is challenging to model the complexity of an ecosystem in a laboratory; however, to gain insight on how ecosystems are impacted by outside perturbations it is valuable to develop approaches that mimic an environmental system. Here, we developed a mesocosm that uses readily accessible components that come together to model a terrestrial ecosystem which is coupled with an analysis pipeline to assess how various perturbations impact the soil. We demonstrate the utility of this approach by tracking the effects of three perturbations (water, nutrition, contamination with yeast) on the soil over the course of 30 days. Our results demonstrate that these treatments can have lasting impacts on the soil. These findings and the methods presented here could be useful to other researchers assessing how ecosystems respond to perturbations.

**Highlights:**

- We developed a pipeline using terrestrial mesocosms that allow for the analysis of how perturbations impact soil systems and demonstrate that it is effective for targeted detection of a microbe of interest as well as global phylogenomic observation of ecological changes due to external perturbation events.
- digital droplet PCR was adapted to track a low abundance, non-native microbe in soil mesocosms.
- Temporal sampling allowed for the longitudinal observation of soil response to a one-time perturbance.
- Introduction of yeast and its associated growth media conferred an expansion of total biomass and increase in alpha-diversity and shifts in the beta-diversity of the soil microbiome.
- Treatment with media or yeast resulted in the expansion in the relative contribution of fungal biomass and an increase in the relative abundances of *Saccharomycetes and Trellomycetes,* with decreases in *Sordariomycetes, Leotiomycetes, and Eurotomycetes*
- Media or yeast *introduction* also resulted in an expansion of the relative abundances of *Gammaproteobacteria, Bacilli,* and *Bacteroidia,* and decreases in *Actinomycetia* and *Acidobacteria*.

## Introduction

Soils perform many critical processes that are fundamental to ecosystem function and planetary health and serve as a key global carbon sink^1^. Soils host dynamic and complex ecosystems, teeming with microbial life that are crucial to ecosystem function, plant productivity and consequently, societal and human health^2,3^. Globally, soils are eroding and degrading at alarming rates due to increased perturbances driven in large part by land use and climate change events^4^. Perturbances to soils can have long lasting and wide-reaching impacts, such as reduced plant productivity and decreased carbon capture capacity ^5^. Thus, modeling soil perturbations has broad utility, from predicting the impacts of climate-related events or agricultural practices, to developing and validating biobased agronomy strategies or bioremediation efforts. And, of particular interest to our work, such approaches enable the assessment of potential environmental impacts of industrial biotechnologies and the introduction of non-native microbial strains upon soil ecology. The assessment of the impacts of perturbation events on soil is warranted, as research on disturbances such as wildfires, heatwaves, and drought have indicated that a one-time perturbance can have widespread and sustained impacts on soils ^6–8^, however modeling a soil ecosystem in a streamlined, high-throughput and reproducible manner in a laboratory environment can be challenging, therefore we developed an accessible mesocosm-based system and analysis pipeline that would allow for the evaluation of how various perturbations or contamination events would impact a soil system. Here we present the findings from using this platform to evaluate the impact of three perturbation events: hydration (water amendment), nutrification (media amendment), and contamination, via introduction of the industrial microbe, *Saccharomyces cerevisiae,* however this simple system could be readily adapted to probe myriad questions on the impacts of perturbations, amendments, or contamination events upon soils and associated plants. Yeast was selected because it is an industrially relevant microbe that is important to multiple fields, including the production of biofuels ^9–12^. It is important to note that the practical application of these technologies at scale may necessitate genetic modification of *S. cerevisiae*, despite the fact that the potential environmental impact of such modifications, or even of a large scale contamination event with wild-type yeast, remain poorly understood.^13–15^. To gain insight on the potential impacts of a soil perturbation we sought to develop a lab-scale terrestrial mesocosm that could be used as a platform to assess the impact of various perturbations on both the abiotic properties of the soil and the underlying microbial communities.

Here, we describe a platform for the longitudinal analysis of soil microbial communities that utilizes mesocosms coupled with experimental and informatic analyses to assess the effects of three types of perturbations upon terrestrial soil and soil microbiome dynamics. Our aim was to develop a flexible and accessible platform that would represent some of the complexity of a terrestrial ecosystem but would be controlled and reproducible enough that it could be used to ask targeted questions and systematically evaluate the impact of environmental perturbation events. Our mesocosms contained potting soil and a *Sorghum bicolor* seedling. Samples were collected over a 30-day time-course following a single “soil perturbance” event. The methodology utilized here allows for minimally disruptive soil sampling over the course of 30-days, assessment of abiotic soil properties at baseline and completion, and time course observations of microbiome dynamics as well as the targeted detection of a microbe of interest in a dense and complex microbial ecosystem. The resultant data support that: (1) our mesocosms host a rich ecosystem that is dynamic and multitrophic, (2) our mesocosms have strong agreement between replicates, (3) our approach allows for the broad assessment of shifts in the microbial community, and (4) that all perturbations had lasting impacts on abiotic soil properties as well as microbial community dynamics. We were also able to track the fate of yeast over time in the soil, and although it failed to survive in the mesocosms it had lasting impacts on the underlying microbial community.

These data not only shed light on how a perturbation event may impact the dynamics of a soil ecosystem and associated microbiome over time, but it also serves as a facile methodological framework for a lab-based assessment of soil perturbation events and the specific effects of the introduction of a microbe of industrial importance on the soil ecosystem. These findings underscore the efficacy of this approach for assessing soil perturbances and support the argument that risk assessments of industrial microbes are warranted.

## 1. Results

### Abiotic soil properties were altered following perturbations

Mesocosms were assessed over a 30-day time-course following a single “soil perturbance” event, which entailed treatment with water, YPD media, or yeast in YPD-media (**Fig. 1)**. The yeast strain used in this study constitutively expressed GFP to facilitate fluorescent microscopic detection. Abiotic soil properties, including carbon and macronutrient concentrations, were evaluated at baseline (t_0_) and at the culmination of the experiment (t_30days_) to establish the impact of perturbances.

**Figure 1.**
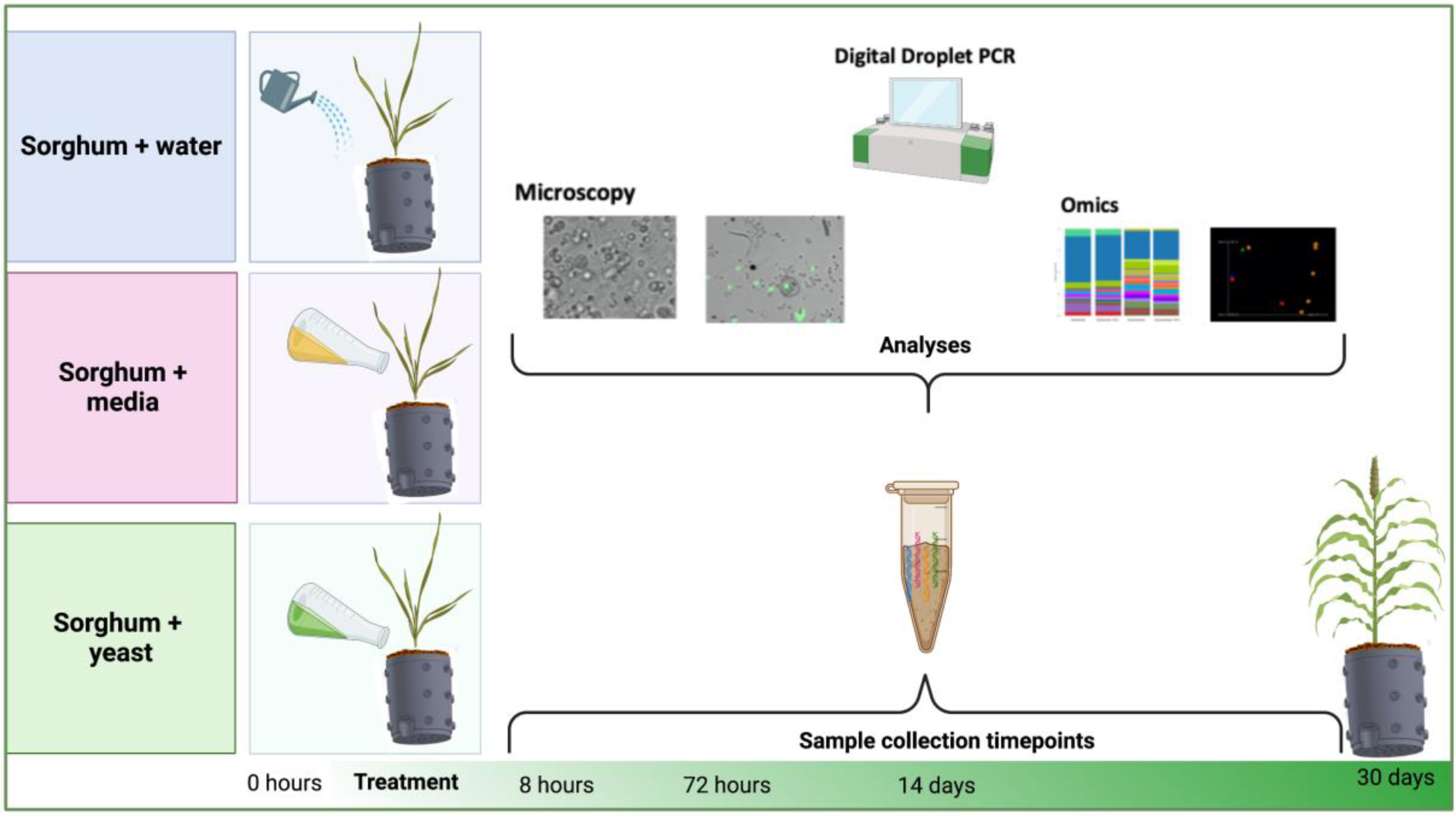
Experimental Design. Samples were collected from mesocosms immediately before treatment at baseline (0 hours), then subsequently at 8 hours, 72 hours, 14- and 30-days post treatment. All samples were observed with light microscopy and extracted genomic DNA was used for targeted 16S and ITS sequencing and digital droplet PCR. Bulk soil was assessed with PFLA and Haney testing at baseline and 30 days. Schematic created with BioRender.com

Inorganic nitrogen, as measured by H3A extraction, was significantly depleted over the course of 30 days in pots that contained just soil but did not have a S. *bicolor* seedling and in the plants that received water treatment **(Fig. 2A**). This depletion was largely mitigated in soil that received either media or yeast; ammonium was increased in both media treatment and yeast treatment (**Fig. 2B**). Phosphorus followed a similar pattern of depletion, however, only mesocosms that received yeast had insignificant depletion of phosphorus at endpoint (**Fig. 2C**). No significant changes in soil pH were observed at any time point or with any treatment (data not shown). Percent microbial active carbon (%MAC) was moderately reduced in yeast treatment (<10%). Soil respiration (CO_2_-C) and Soil Health Calculation (1 day CO2-C divided by organic C:N ratio plus a weighted organic carbon and organic N addition) were significantly increased in plants that received media or yeast (**Fig. 2D**).

**Figure 2.**
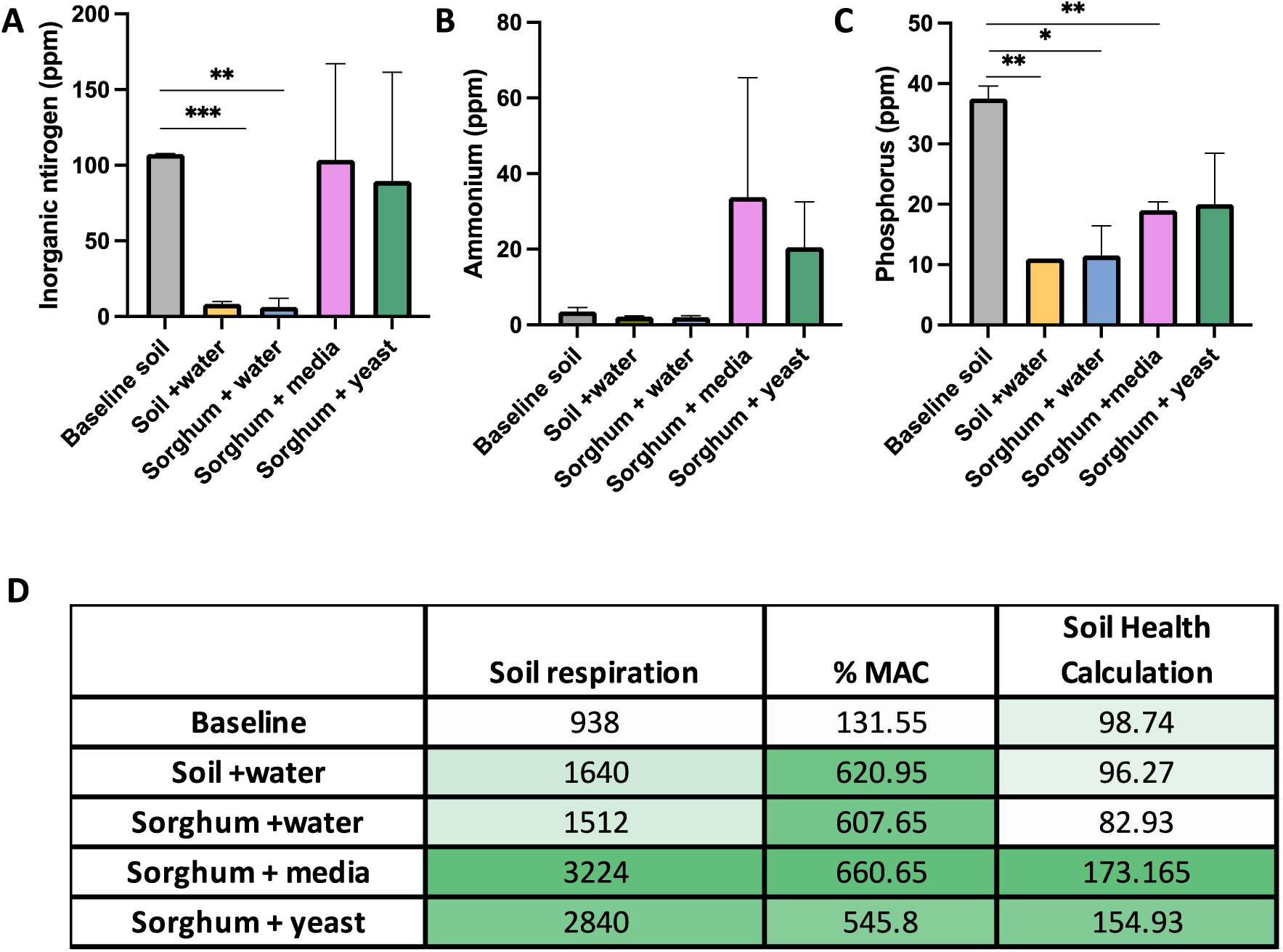
Haney Analysis of Soil. Changes in nutrients by treatment at the 30-day timepoint versus baseline as evaluated by HA3-1 soil extractant. Inorganic nitrogen (**A**), ammonia (**B**) and phosphorus (**C**). Table of soil respiration, % microbially active carbon, and soil health calculation by treatment at 30 days versus baseline(**D**).

### The impacts of amendments on the biological composition of the soil were detectable up to 30 days after perturbations

Phospholipid fatty acid (PFLA) analysis was used to assess the broad phylogenetic microbial composition of soils at baseline and 30 days after perturbation (+water, +media, or +yeast). Microbial biomass totals increased with all perturbations over the course of 30 days, with the most pronounced increases being in the samples that received yeast (**Fig. 3A)**. In addition to increased biomass, the relative contributions of functional groups shifted with soil perturbations, with the percent of fungal biomass increasing in samples that received media or yeast versus samples that only received water (26.02% vs. 13.01, p-value=0.02; 23.75% vs. 13.01% p-value 0.05). These changes are mirrored in the fungal: bacterial ratios **(S. Fig. 2),** which were shifted toward a greater fungal contribution in samples that received media and were significantly increased in yeast treated samples (p=0.04).

**Figure 3.**
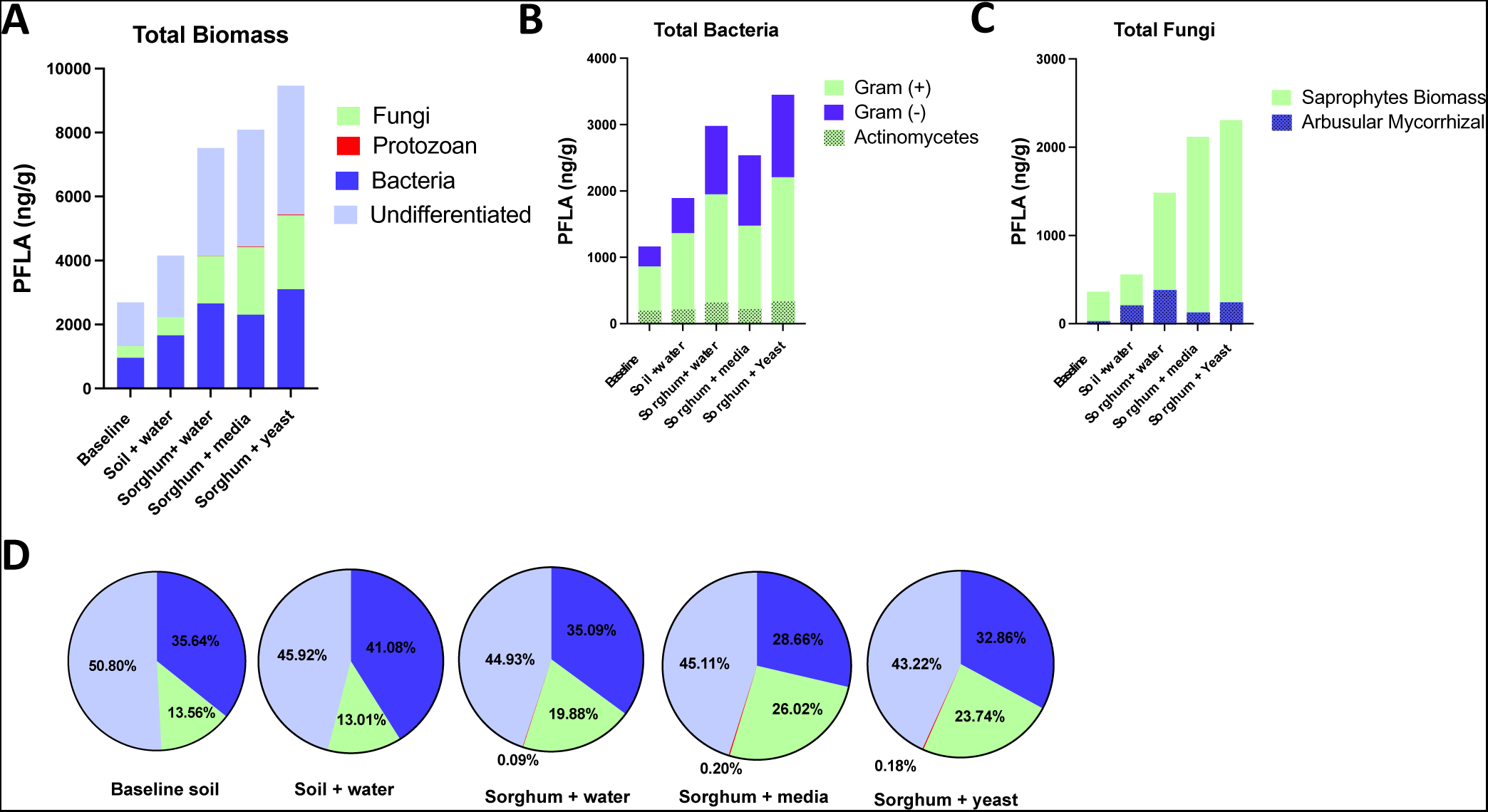
Phospholipid fatty acid (PFLA) analysis by Gas Chromatography (GC). Microbial composition of soils at baseline and 30 days post amendment. **A)** Total biomass (sum of PFLAs) **B**) breakdown of bacterial and fungal biomass **C)** Percent composition of biomass. Results represent the average of biological duplicates of each sample.

### The phylogenetic diversity of the soil microbiomes increases over time and as a function of perturbation events

Targeted sequencing of the 16S and ITS genes was used to compare the bacterial and fungal communities of soil samples over the duration of the experiment. Bacterial and fungal communities shifted over the 30-day time course regardless of perturbation with both the phylogenetic diversity of individual samples (alpha diversity) and the community composition of samples (beta diversity) shifting over time (**S. Fig. 3)**. The mesocosms also resulted in stratified communities with samples collected from the upper ports being more impacted by treatment than samples collected from the lower ports (**S. Fig 4)**. Impacts of the perturbations were more pronounced in the upper strata; thus, subsequent results focus on the top ports only.

The alpha diversity, or phylogenetic diversity of the microbes in a given sample, was increased in all groups over time, but mesocosms that were perturbed via nutrification (media), or contamination (yeast) had higher alpha diversity of both the fungal (F) and bacterial (B) components of the soils, as measured by Faith’s phylogenetic diversity (PD). Across all samples, alpha-diversity increased as a function of time, regardless of perturbation, with all groups having significantly higher Faith values at the 30-day time point versus earlier timepoints; 0 hours (F&B p = <0.001), 8 hours (F &B p= 0.001), and 72 hours (F p= 0.01; B p= <0.001). However, there were additional increases in alpha-diversity driven by the perturbations themselves. The highest alpha diversity was in the yeast group at 30 days, with the delta between day 0 and day 30 being greater than that of media or water treatment (Fungal; 11.56 vs. 8.48 vs 5.69, Bacterial; 52.91 vs. 20.04 vs 28.48). (**Fig. 4)**

**Figure 4.**
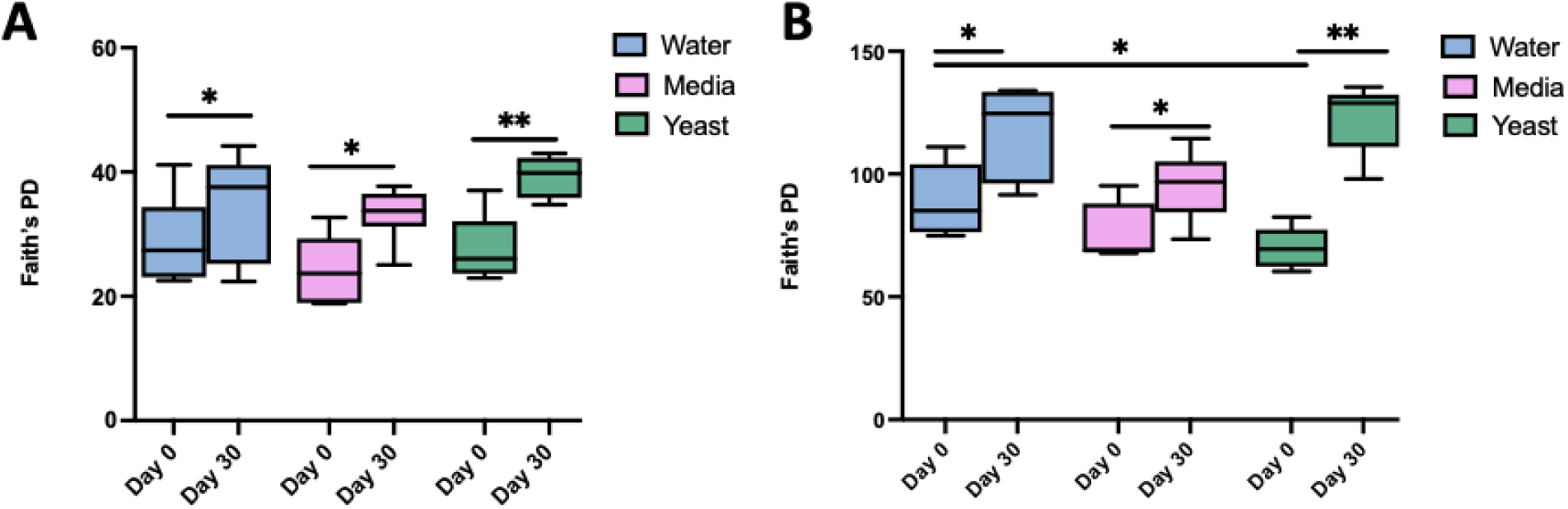
Alpha diversity indices of soil microbiome increase over time and as a function of perturbation. Alpha diversity of the fungal (**A**) and bacterial (**B**) soil sample as a function of perturbation at a given time point as assessed by Faith’s PD which assesses phylogenetic units in a sample.

### Community composition of soil microbes is differentially shifted as a function of perturbation

The beta diversity, or overall composition of the soil microbial communities, was shifted following perturbations. To assess how each perturbation differentially impacted the soil microbiome we compared beta-diversities between treatment groups. Beta diversity shifted both over time and also as a result of each perturbation, indicating that perturbation type uniquely impacted the overall microbial composition of the soils compared to baseline (**Fig. 5**).

**Figure 5.**
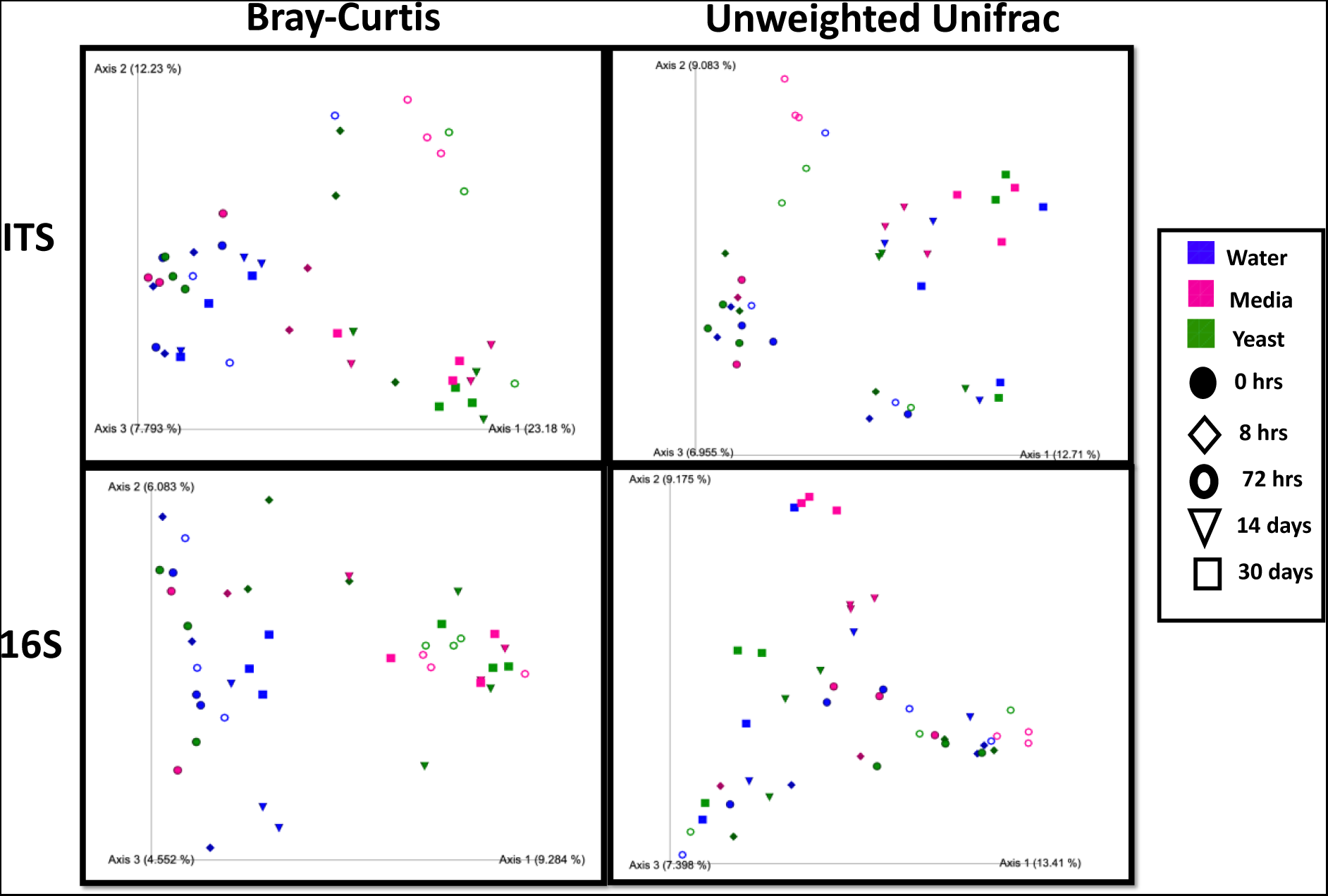
Beta diversity indices of soil microbiome increases over time and as a function of perturbation. Perturbations shift the beta-diversity of soil mycobiome (top) and bacteriome (bottom) with Bray-Curtis distance indices (left) and Unweighted Unifrac distance indices (right) with time indicated by icon shape and treatment by icon color. Fungal (top) and bacterial (bottom) diversity of soil samples by time and treatment. Statistical significance of timepoint or treatment assessed by multivariate regression analysis Adonis; model = timepoint + treatment.

Media or yeast significantly altered the fungal community of the soil compared to water when evaluated with beta-diversity metrics (Bray-Curtis, Unweighted Unifrac, or Weighted Unifrac; Pairwise PERMANOVA <0.01). At the 30-day timepoint, the distance between water treated soils and yeast treatment was greater than that between media and water using (Bray-Curtis, Unweighted Unifrac, or Weighted Unifrac) (**Fig. 5).** However, the distance between media treated and yeast treated soils were only significantly different by Unweighted Unifrac (p = 0.004). Analysis using the Adonis model showed significance of both timepoint and treatment independent from one another with all p-values being ≤ .001.

The bacterial communities also exhibited significant shifts in composition as a function of time and of treatment with both media and yeast mesocosms being significantly diverged from water treatment as well as from each other at most timepoints, as assessed by beta-diversity metrics (Bray-Curtis, Unweighted Unifrac, or Weighted Unifrac), with one exception being there were insignificant differences between water and yeast at 30 days as measured by Unweighted Unifrac, which is unsurprising in that all mesocosms started with fairly uniform communities and experiments were carried out in a closed system, suggesting that most changes were driven by shifts in relative abundances and expansions of low level taxa. There were no significant differences in beta-diversity among the baseline samples (as measured by any of the above metrics) indicating homogeneity between mesocosms (**S. Fig. 4).**

### Different classes in fungal and bacterial communities became dominant following perturbation events with yeast driving the greatest shifts

The dominant fungal classes were shifted over time as a function of perturbation with the *Sacchromycetes* increasing in both the media (p = .009) and Yeast (p = .001) treated soils versus water treated, *Tremellomycetes* were similarly increased in media (p = .04) and Yeast (p = .008) with a concurrent reductions in *Sordariomycetes* (media p = .04, yeast p *=* .006), *Leotiomycetes* (media p = .0002, yeast p = .001), and *Eurotiomycetes* (media p = .01, yeast p = .003) (**Fig 6A**). To assess if the expansion in *Saccharomycetes* in the soils that received yeast was driven by a proliferation of yeast itself, we homed in on this group. However, the genus *Saccharomyces* was only detected with ITS sequencing at the 8-hour time point, where it accounted for an average of 23.7% relative abundance at the genus level in the plants that received the yeast amendment (data not shown). We next deployed targeted detection methods to assess the persistence and fate of yeast using fluorescence microscopy enabled by constitutive GFP expression. Yeast was observed in soil samples up to 72 hours after treatment (**Fig. 6A**) but was not seen at later time points and could not be quantified. Using digital droplet PCR (ddPCR) we were able to detect and quantify yeast genomic DNA from soil extractions for up to 14 days post treatment using a probe specific to the *spa2* gene^16^ (**Fig. 6B**).

**Figure 6.**
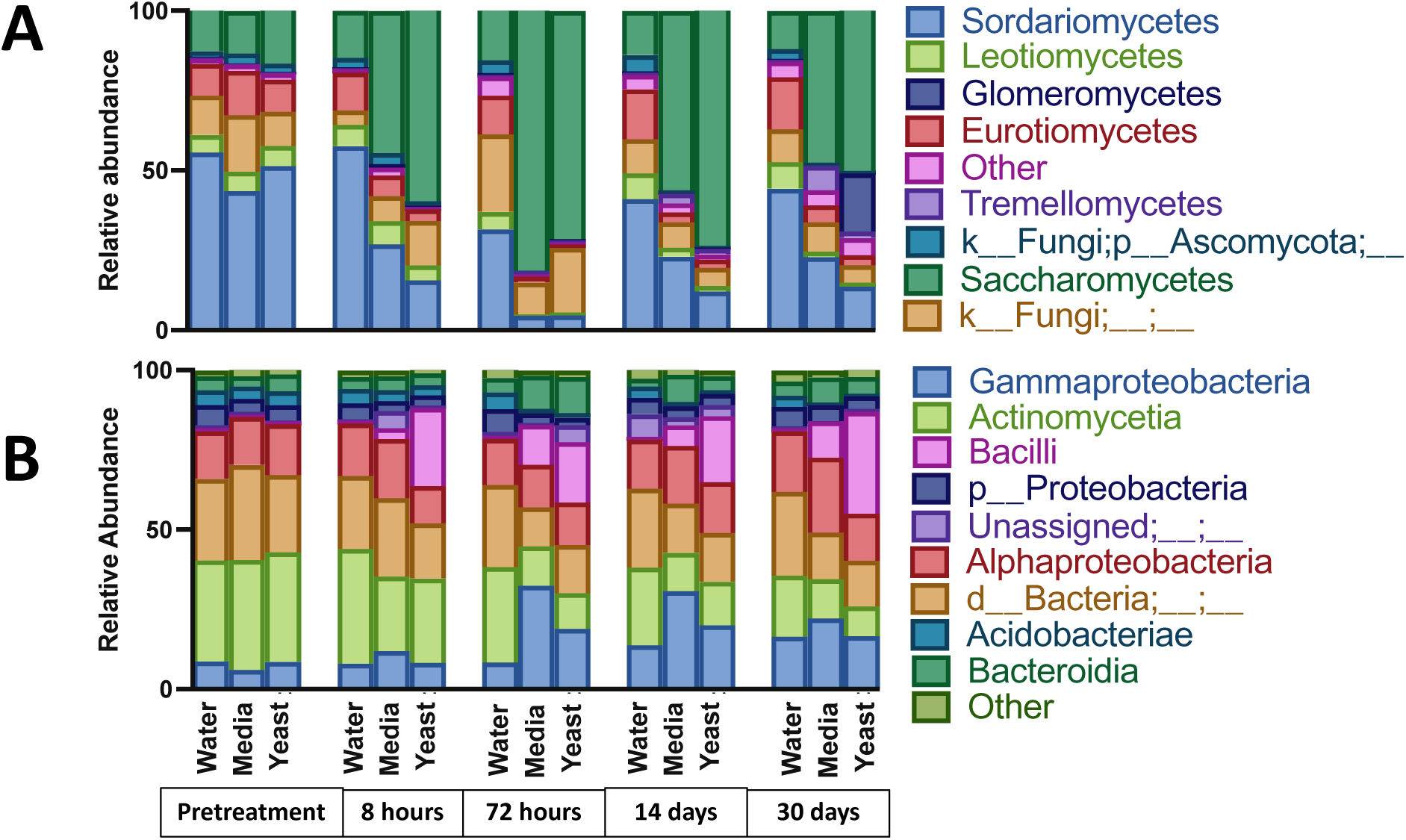
Taxonomic profiles of dominant classes over time and by treatment. Taxa bar charts showing relative abundance of fungal (A) and bacterial (B) classes by time point and treatment. Legend identifies the topmost abundant classes (Contributing > 5% Relative Abundance.)

**Figure 7.**
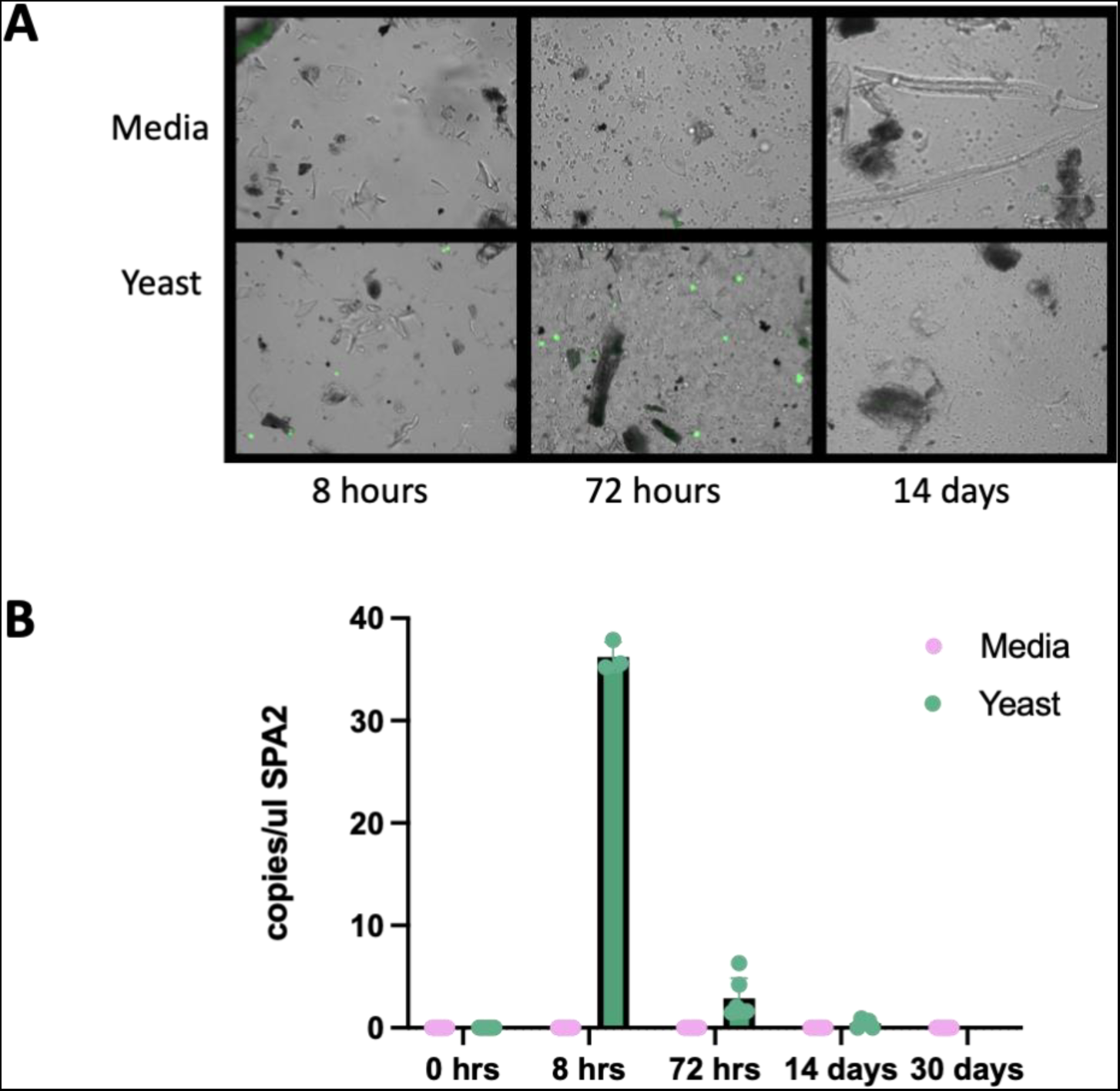
Direct detection of yeast in mesocosm over time. **A)** Fluorescence images of soil aliquots at various time points, yeast in green, all images taken at 40X. Images are representative of slides prepared in technical triplicate from biological triplicates. **B)** Detection of yeast using digital droplet PCR on total soil gDNA targeting the *spa2* gene.

Shifts in the bacterial community included a significant expansion of the class *Bacilli,* the relative abundance of which was increased in both media and yeast vs. water but to a significantly greater extent in yeast treated plants (water vs. yeast p = .001; water vs. media p = .01; media vs. yeast p = .01). Increases in *Gammaproteobacteria* and *Bacteroida* were also observed. Both media and yeast also conferred increases in *Gamaproteobacteria*, and *Bacteroidia*. These increases coincided with reductions in *Actinomycetes*, *Proteobacteria* and *Acidobacteria*. (**Fig. 6B**).

## Discussion

Rapid advancements in the development and application of industrial microbes are paving the way towards the realization of a bioeconomy; however, the potential risks these microbes may pose has yet to be thoroughly evaluated. This is due, in part, to the challenge of modeling inherently complex and dynamic environmental conditions under laboratory constraints. Despite these challenges, a systematic assessment of environmental risks is warranted. Here, we demonstrate the utility of a lab-based soil-sorghum mesocosm to assess the impact of perturbance events on soil characteristics and microbiome dynamics. This approach takes advantage of readily available components (3D-printed pot with sampling ports, commercial potting soil, *S. bicolor* seeds) and results in a dynamic terrestrial ecosystem that allows for minimally disruptive sampling over time, which showed strong reproducibility as well as clear spatial and temporal resolution. This approach could thus be useful for many types of studies; here, we chose to monitor the impacts of soil perturbation events. While there are myriad perturbations that could alter soil characteristics, here we used the system to assess the impact of three perturbance events; hydration (water treatment), nutrification (media treatment), or contamination (inoculation with *S. cerevisiae*). This approach allowed us to evaluate the impact of a perturbation on the underlying soil microbiome over time as well as assess the survival and fate of yeast in the soil ecosystem.

We composed our mesocosms of commercial potting soil and *Sorghum bicolor* seedlings which were maintained in a plant growth chamber for 30 days. Soil was chosen as it is critical to ecosystem health and productivity and introduces high levels of microbial diversity ^2,17,18^. The commercial potting soil used here introduced the microbial complexity needed to model an environmental escape but was uniform enough that it could be used for replicates and validation. Our mesocosms also contained a *S. bicolor* seedling, as plants are critical to maturing and stabilizing soil microbiomes^19^. Additionally, *Sorghum spp,* has significant potential both as a food crop and a bioenergy feedstock, making it particularly relevant to our evaluation of an industrial microbe^20,21^.

Analysis of our soils by PFLA and Haney testing indicate that we have a dynamic soil system that is trophically diverse, effectively modeling a true phytobiome in which we could assess the systematic impacts of soil perturbations. Importantly, our system showed strong agreement between baseline samples, as well as within treatment groups, supporting that it captures the complexity of a multi-trophic soil system but is still reproducible and stable. All perturbations increased soil biomass, nutrification with media amendment and to a greater extent contamination with yeast both increased fungal representation. Nutrification or yeast supplementation resulted in a higher fungal: bacterial ratio which canonically has been considered indicative of soil health in agricultural soils ^22^.

The yeast strain assessed here failed to survive and was only detectable for 72 hours via microscopy with genetic signatures being detected for up to two weeks using ddPCR. Direct detection of a low abundance microbe in the crowded microbial world of soil is a unique challenge, as classical techniques such as plating on permissive or selective media are futile in the face of metabolically diverse and antibiotic resistant soil microbes. Microscopic observation is useful for qualitative observation but cannot quantitively capture the entirety of the system. Traditional PCR can be challenging in soil samples as the sheer volume of templates as well as an abundance of PCR inhibitors, such as humic acid, can interfere with PCR chemistry ^23^. Here, we demonstrate that ddPCR can overcome some of the barriers to detection of low abundance targets in soil and proved highly sensitive, making it a powerful option for analyses of this nature. Future analyses will evaluate if the DNA detected with ddPCR at later time points originated from live yeast that was persisting in the soil versus relic DNA from dead yeast, which has been noted as an issue with sequencing data from soils ^24^.

Despite the transient survival of yeast in the soils, the impact of a one-time perturbation was persistent, having effects on the soil microbiome that were greater than those caused by hydration or nutrification for the full 30 days of observation. All perturbations led to increased alpha-diversity and sustained differences in beta-diversity. Increases in alpha-diversity were likely driven by expansions of low-abundance taxa, as the soils were largely homogeneous in composition at the onset. Increased alpha-diversity is canonically associated with heartier and more resilient soil systems ^25,26,27^, thus it would be compelling to extend these experiments to see if the increased alpha-diversity is maintained beyond 30 days and if it has any effects on plant productivity. Future studies will also explore if increased diversity indeed enhances the resilience of the soils to subsequent perturbations.

The shifts in community composition across treatments were perhaps the most compelling results of the study; though all soils matured over time, resulting in changes to beta-diversity, the soils that received media or yeast underwent the most profound shifts. We hypothesize that the sustained expansion of the *Sacchromycetes* in these treatment groups was largely driven by readily available food sources that the yeast extract in YPD media and the yeast itself provides, as the *Sacchrmoycetes* are saprophytes. Future analyses will elucidate if the nutrients associated with different perturbations could predict the expansion of taxonomic groups, i.e., would a nutrient influx that favored a specific bacteria result in greater blooms of that bacterial taxa? Intriguingly, plants that received yeast supplementation also demonstrated an expansion in the *Tremellomycetes* at 30 days, which represent important members of the rhizosphere^28^ .

The bacterial component of the soils was also significantly impacted by the perturbations with an expansion of unclassified bacteria in the plants that received yeast that coincided with a reduction in both the *Acidobacteria* and the *Actinobacteria*, both of which are major players in soil microbial ecology with myriad ecological niches and functional activities ^29,30^. Extension of these experiments would not only allow us to evaluate if said changes in microbial composition have functional impact, but it would also allow us to assess if the soils ultimately rebound from the perturbance, or if the change in the trajectory of community composition is sustained.

These data show that this in-lab mesocosm pipeline can be used to comprehensively evaluate the potential impact a microbial perturbance event may have in a soil model that encompasses the complexity and dynamic nature of terrestrial soils. Further, this system is useful for assessing the risk of persistence of an exogenous microbe in the environment, and to gauge the long-term impact and soil rebound capacity following perturbation. The profound and prolonged shifts that resulted from either media treatment or yeast treatment, despite the short survival of yeast in the mesocosms, underscore the importance of evaluating potential impacts of industrial microbes and their associated growth media and product suites on native microbial ecologies to avoid unwanted disruptions to ecosystems, such that we can responsibly progress in the development and deployment of industrial microbes to advance the realization of a sustainable and productive bioeconomy.

## Methods

### Yeast strain development and cultivation

*Saccharomyces cerevisiae* BY4741-derivative strain GabY36 (*MATa his3Δ1 leu2Δ0 met15Δ0 ura3Δ0 XI-3: CamPr-GFP XII-5: TDH3-Cam-TA*) was transformed with plasmid GabP54 [pCfB2904 (XI-3 MarkerFree), with the *CamP* driving expression of GFP] was transformed to contain a GFP expression cassette^31^. pCfB2904(XI-3 MarkerFree) was a gift from Irina Borodina (Addgene plasmid # 73276; http://n2t.net/addgene:73276 ; RRID:Addgene_73276) General yeast manipulation, media, and transformation were performed by standard methods (Amberg, 2000). For CRISPR mediated construction, protocol was followed as described previously^32^. Yeast was maintained in YPD +20% glucose (yeast extract (10 g L^−1^), peptone (20 g L^−1^) and glucose (20 g L^−1^); agar (20 g L^−1^) was used for plates.

### Mesocosm experimental design and sampling procedure

*Sorghum bicolor* BTx623 seeds (shared by Scott Sattler USDA-^29,30^) were started in seedling trays containing ∼25 ml packed commercial potting soil *Soil Sunshine Mix #4* (Sungro, Agawam, MA) and cultivated in a Percival Growth Chamber E-41VL (Percival Scientific, Perry IA) at 25°C, under diurnal light conditions for 1 month at which point the seedlings reached five leaf coleoptile developmental stage.

Mesocosm experiments were conducted in 3D-printed pots (**S. Fig. 1)** that have 18 sampling ports which are sealed with neoprene stoppers (Eisco Labs, Victor, NY). A perforated base allows for root development. Plants self-water via wicking from a reservoir (Nalgene, Rochester, NY). To facilitate self-watering prior to root establishment each pot has a length of capillary wick chord from the pot to the reservoir (ORIMERC, USA).

*Soil Sunshine Mix #4* was thoroughly mixed for homogenization and moistened with DI water, each pot was firmly packed with damp soil, and sorghum seedlings were transplanted to mesocosm pots and allowed to settle for 72 hours under the above growth conditions to stabilize the microbial community.

72 hours post-transplant, baseline samples were harvested from one set of ports on each plant. Briefly, a 3 ml syringe (BD Biosciences, Franklin Lakes, NJ) was modified by cleaving the top off and was then used to extract a horizontal soil core from each port. Soil samples were ∼ 250 mg and either went directly into the lysis buffer for gDNA extraction (as described below) or were resuspended in 800 ml DI water and vortexed to be observed microscopically.

Immediately following the collection of baseline samples, mesocosms were inoculated with one of the following treatments, each was performed in triplicate: **1)** 200 ml of DI water **2)** 200 ml of YPD Yeast -Extract-Peptone-Dextrose (YPD) + 2% Glucose **3)** 200mL of 0.1 OD BY4171 resuspended in fresh Yeast -Extract-Peptone-Dextrose (YPD) + 2% Glucose.

### Microscopy

Soil aliquots were imaged at 40x on a EVOS M5000 (Thermo Fisher Scientific, USA) with trans light and through a GFP filter cube. Slides were scanned and imaged at a minimum of 3 fields of view. Representative images were chosen for each timepoint.

### Plant Observations

All plants were started from seed and experiments were initiated when seedlings reached coleoptile stage (∼30 days from starting seeds). Soil was removed from roots for endpoint analysis and plant matter was air-dried at 25°C and weighed to determine final dry weight.

### Biological composition and diversity and physicochemical characterization of soils

An aliquot of the soil slurry used for the initial setup of the mesocosms and 250 grams of final soil from representative treatment groups were collected and analyzed by Ward Laboratories Inc (Kearney, NB). Community composition was assessed by Phospholipid Fatty Acid (PFLA) analysis. Employing Ward Laboratories standard protocols fresh soil is freeze dried, weighed into flasks, and combined with a 1:2:0.8 (vol:vol:vol) mixture of extracting solvents. The samples shake, then are centrifuged to allow for the separation of the organic fraction. The organic fraction is removed and now contains the fatty acids from the soils. Fatty acids can either be methylated to represent TSFAME or separated into neutral, glycolipids, and phospholipid fatty acids (PLFA) by solid phase extraction (SPE). Following SPE the desired fraction is then methylated. Samples are analyzed on a GC using Agilent’s ChemStation and MIDI’s Sherlock software systems. Soil respiration was assessed as follows; soils are dried and ground by standard laboratory procedures. A plastic beaker with holes drilled in the bottom and a piece of filter paper lining it is used to contain a 40 g (± 0.5 g) sample. The beaker is placed in a half pint mason jar and 20 mL of Deionized Water is added to the bottom of the jar. The jar is sealed and incubated at 24° C for 24 hours (± 1 hour). After incubation, the SR-1 instrument is used to measure the CO2 produced.” Physical and chemical traits of the soil were assessed by Haney Soil Health testing. “Haney Analysis Soils that have been dried and ground by standard laboratory procedures are scooped into appropriate containers in duplicate. One sub sample is extracted with H3A and the other with H2O. The H3A extract is analyzed for NO3, NH4, and PO4 on a Flow Injection Analyzer (FIA) and P, K, Ca, Mg, Na, Zn, Fe, Mn, Cu, S and Al are run on a Thermo ICAP. The H2O extract is also analyzed for PO4 on Flow Injection Analyzer but is also analyzed for Total Organic Carbon (TOC) and Total Nitrogen (TN) using a Teledyne-Tekmar Torch.

### DNA Extraction and Sequencing

Total genomic DNA (gDNA) was extracted from soil samples collected from the top port at each timepoint using the DNeasy PowerSoil Kit (Qiagen, Germantown, MD). Modifications to the standard protocol included a 10-min incubation at 65 °C immediately following the addition of the lysis buffer, cell disruption and homogenization was done with a Digital Cell Disruptor Genie (Scientific Industries, Bohemia, NY) at 3000 RPM for 10 min. Extracted gDNA was normalized to 20ng/ul and sent to Genewiz (Azenta Lifesciences, USA) for targeted sequencing of the 16S and ITS hypervariable regions as follows. Next generation sequencing library preparations, Illumina MiSeq sequencing, and data analysis were conducted at AZENTA, Inc. (South Plainfield, NJ, USA). Employing AZENTAs standard protocol for 16S/ITS-EZ sequencing, the sequencing library was constructed using an ITS-2 Library Preparation kit (AZENTA, Inc., South Plainfield, NJ, USA). Briefly, 50 ng DNA was used to generate amplicons that cover ITS-2 hypervariable region of fungal ribosomal internal transcribed spacer (ITS) region. Indexed adapters were added to the ends of the ITS-2 amplicons by limited cycle PCR. Sequencing libraries were validated using an Agilent 4200 TapeStation (Agilent Technologies, Palo Alto, CA, USA), and quantified by Qubit 2.0 Fluorometer (Invitrogen, Carlsbad, CA) as well as by quantitative PCR (KAPA Biosystems, Wilmington, MA, USA). DNA libraries were multiplexed and loaded on an Illumina MiSeq instrument according to manufacturer’s instructions (Illumina, San Diego, CA, USA). Sequencing was performed using a 2x250 paired end (PE) configuration, image analysis and base calling were conducted by the MiSeq Control Software (MCS) on the MiSeq instrument.” “16S-EZ rDNA next generation sequencing library preparations and Illumina sequencing were conducted at Azenta Life Sciences (South Plainfield, NJ, USA). Sequencing library was prepared using a MetaVxTM 16s rDNA Library Preparation kit (Azenta Life Sciences, South Plainfield, NJ, USA). Briefly, the DNA was used to generate amplicons that cover V3 and V4 hypervariable regions of bacteria and archaea 16S rDNA. Indexed adapters were added to the ends of the 16S rDNA amplicons by limited cycle PCR. DNA libraries were validated and quantified before loading. The pooled DNA libraries were loaded on an Illumina MiSeq instrument according to manufacturer’s instructions (Illumina, San Diego, CA, USA). The samples were sequenced using a 2x 250 paired-end (PE) configuration. Image analysis and base calling were conducted by the Illumina Control Software on the Illumina instrument.”

### Community profiling of the microbiome

Bioinformatics on the soil microbiome were performed in QIIME 2 version 2022.2^33^. Single-end FASTQ files were demultiplexed and quality filtered using the q2-demux plugin and denoising was performed with DADA2^34^. Amplicon sequence variants (ASVs) were aligned with MAFFT ^35^and phylogeny was constructed using fasstree2 ^36^. Faith’s Phylogenetic Diversity metric was used to estimate alpha-diversity ^37^. Beta-diversity was assessed by Unweighted and Weighted UniFrac ^38,39^and Bray-Curtis dissimilarity; Principal Coordinate Analysis (PCoA) were estimated using q2-diversity after samples were rarefied to 4000 (16S) and 2300 (ITS) sequences per sample. The q2-feature-classifier, classify-sklearn naive Bayes classifier trained on the Greengenes2 taxonomic database^40^ was used to assign taxonomy to ASVs ^41^ from the bacterial sequences and for fungal sequences we used the UNITE database (V8 Dynamic 10.05.2021 97, 99) ^42,43^. Statistical analyses of alpha-diversity were assessed with pairwise Kruskal-Wallis test, significance of is beta-diversity was assessed with pairwise comparisons using PERMANOVA. To evaluate the individual impacts of variables we employed the ADONIS regression model framework. Changes in relative abundances of microbial classes were evaluated with a Two-War ANOVA, using an interaction term and Geisser-Greenhouse correction.

### Digital Droplet PCR

ddPCR was performed as previously described^44^using the BioRad QX200 with AutoDG system (Bio-Rad, Hercules CA) to monitor the persistence and fate of yeast in soils. A custom HEX fluorophore probe (Bio-Rad) targeting the SPA2 gene of *S. cerevisiae* ^45^ was used on gDNA extracted from total soil **Fwd:** AGAAAACCTTCAGGAACGGG **Internal Oligo (HEX):** CGCCCATAAAGGCAGTAACATCGGC **Rev:** TTTGGCTTGTGGAGGTAGTG. gDNA templates were normalized to 500 pg/rxn and final 20 ul reactions consisted of 2x ddPCR Supermix for Probes (no dUTP) (Bio-Rad), 20x target primer/probes, and DNAse free water. Droplets were produced by AutoDG (BioRad). The sample plate was heat-sealed on a PX1 PCR plate Sealer (BioRad) and thermocycling was performed on a C1000 Touch with a Deep Well Reaction Module (BioRad) using the following conditions: 95°C, 10 min; 94°C 30 sec; 59.4°C, 1:00 min, Ramp 2°C/s, repeat 39x; 98°C, 10 min; 12°C infinite hold. Final droplets were read by the QX2000 Digital PCR Reader (BioRad) and data was analyzed with QuantaSoft analysis package (BioRad) and graphed with GraphPad Software (Dotmatics, San Diego, CA).

## Data Availability

All sequencing data and associated metadata will be publicly available on the National Microbiome Data Collaborative (NMDC) Portal.

## Acknowledgements

We would like to thank Dr. Scott Sattler (USDA-ARS) for providing the Sorghum seeds used in this study as well as useful input on plant growth parameters and Dr. Catherine Lozupone for insightful guidance on study design and manuscript input, and Michael McCausey for assistance with pot design and execution. This work was authored by the National Renewable Energy Laboratory, operated by Alliance for Sustainable Energy, LLC for the U. S. Department of Energy. This work was supported by the U.S. Department of Energy, Office of Science, Office of Biological and Environmental Research, Genomic Science Program under Secure Biosystems Design Science Focus Area IMAGINE BioSecurity: Integrative Modeling and Genome-scale Engineering for Biosystems Security under contract no. DE-AC36-08GO28308. The views expressed in the article do not necessarily represent the views of the DOE or the U.S. Government. The U.S. Government retains and the publisher, by accepting the article for publication, acknowledges that the U.S. Government retains a nonexclusive, paid-up, irrevocable, worldwide license to publish or reproduce the published form of this work, or allow others to do so, for U.S. Government purposes.

---

**Supplemental Figure 1.**
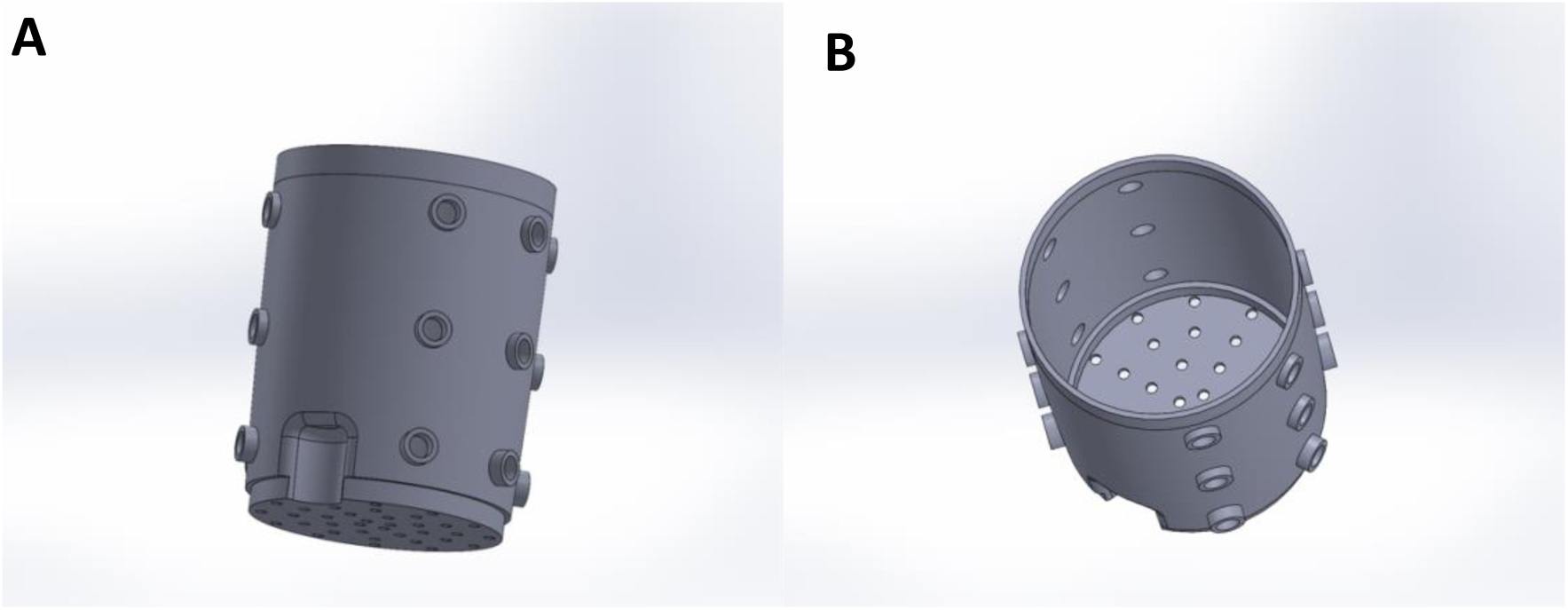
CAD drawing of 3D printed pots showing sampling ports along the side A) side view and B) top view the perforated bottom that allows for root formation and self-watering from a reservoir.

**Supplemental Figure 2.**
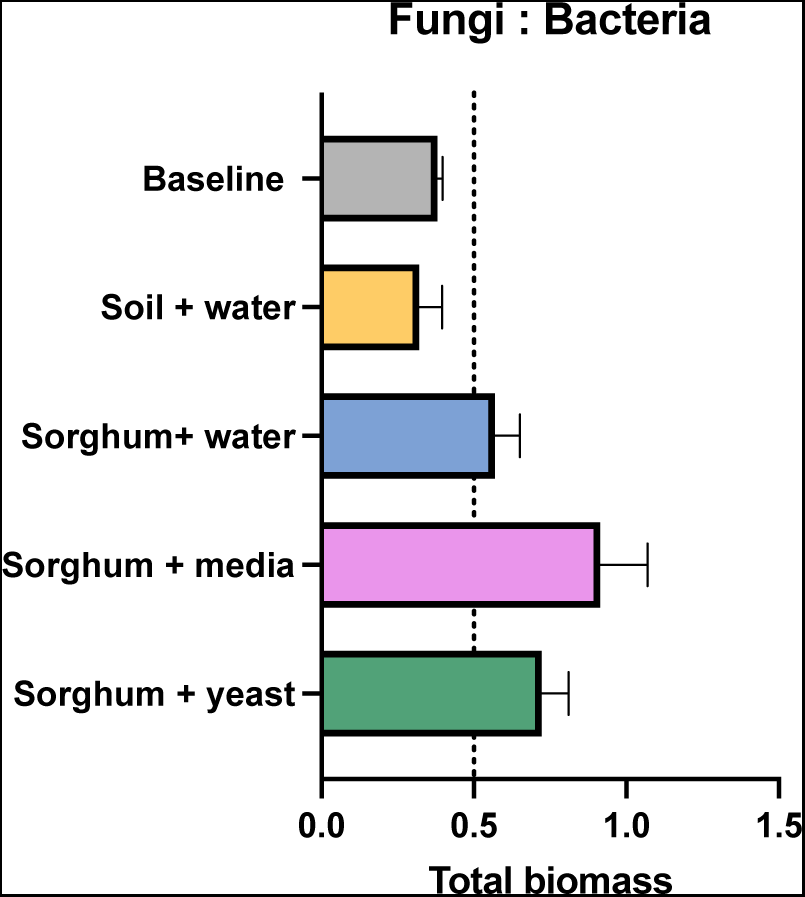
Ratios of fungal biomass to bacterial biomass by phospholipid fatty acid (PFLA) analysis by Gas Chromatography (GC) of microbial composition of soils at baseline and 30 days post amendment.

**Supplemental Figure 3.**
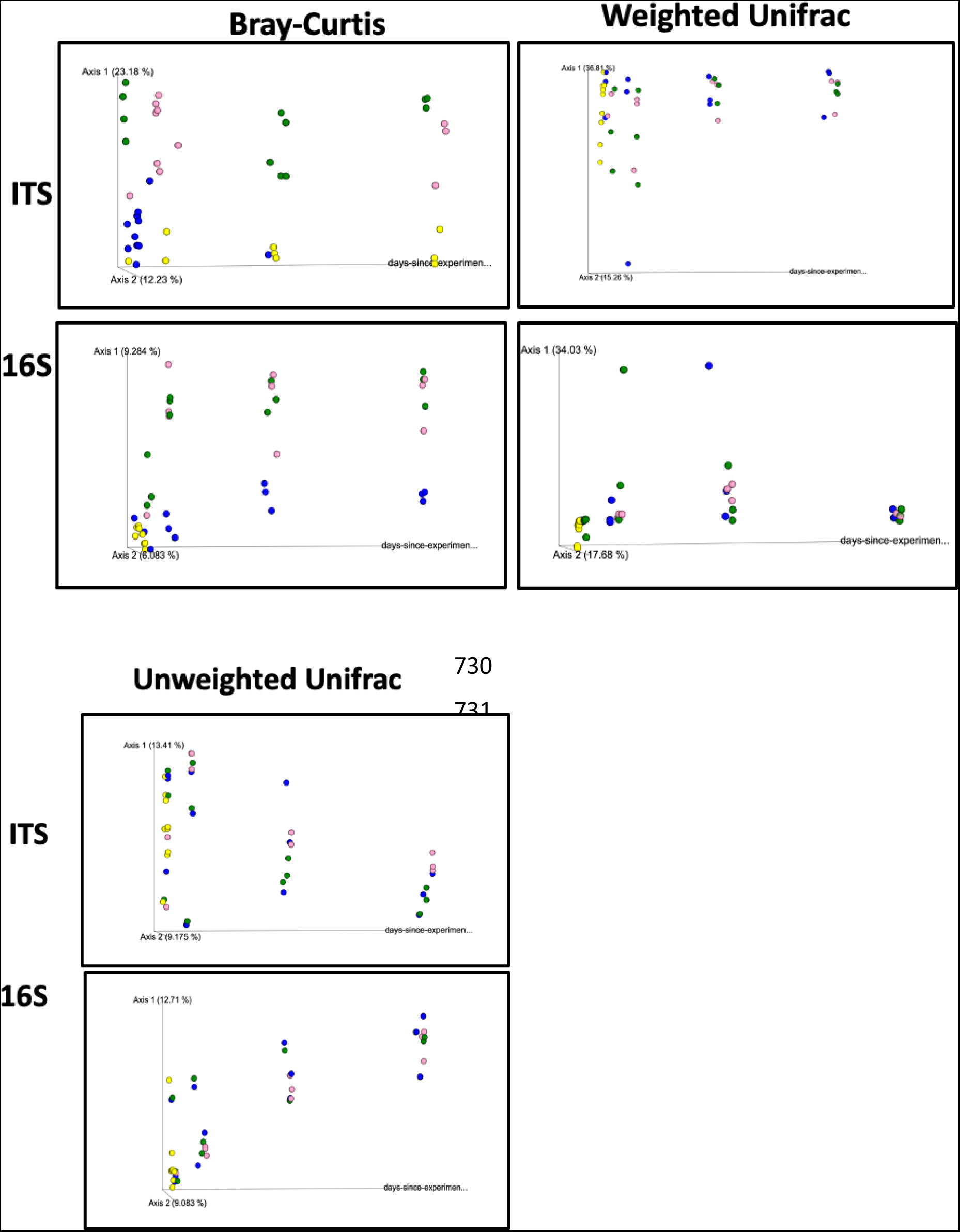
Beta-diversity of samples over time course. Principal coordinate 1, principal coordinate 2, and days since the experiment start to visualize how these samples changed over time.

**Supplemental Figure 4.**
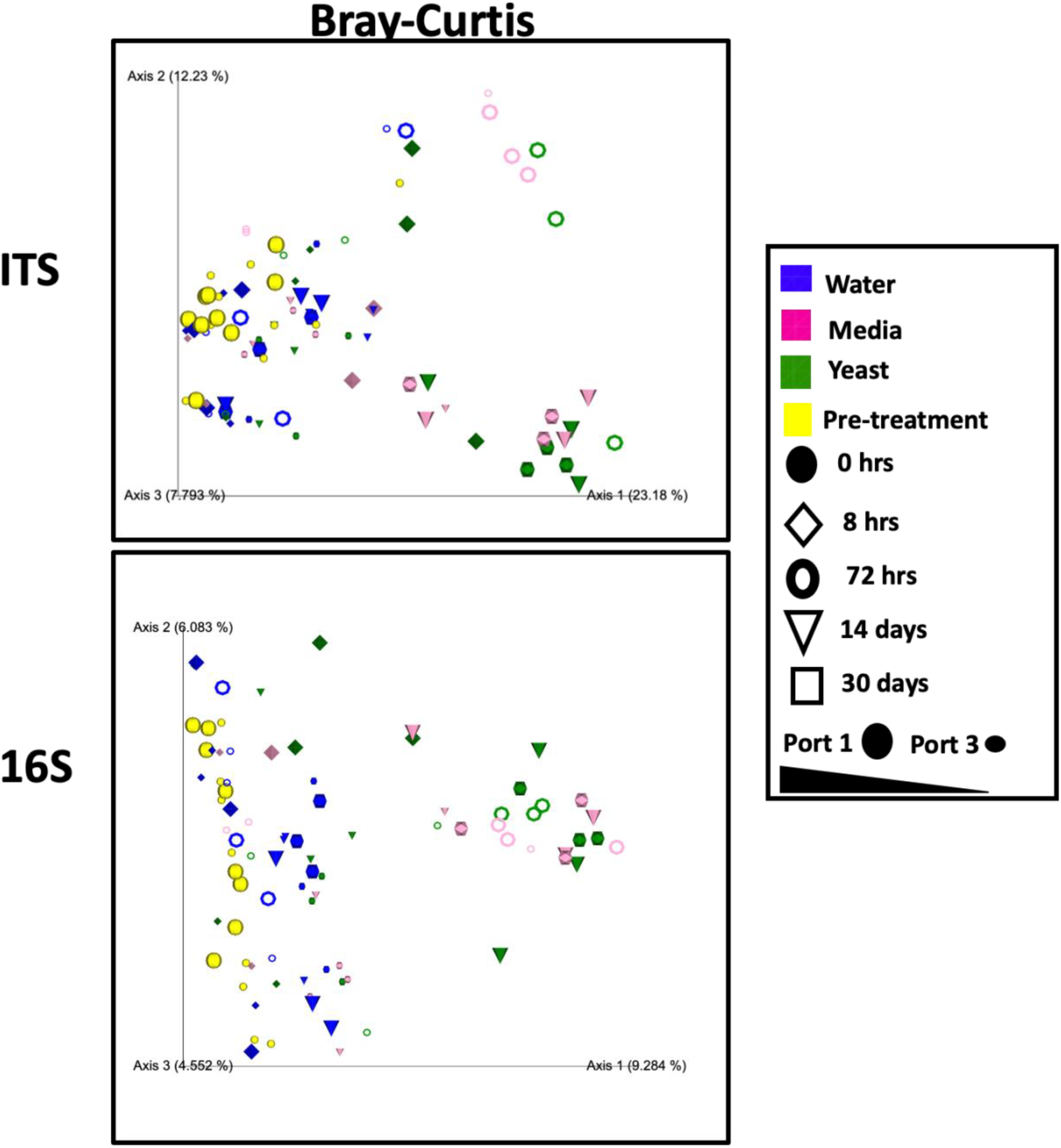
Beta-diversity as a function of treatment (color) time (shape) and strata (scale) of soil mycobiome (top) and bacteriome (bottom) with Bray-Curtis distance matrix.

